# EMReady2: improvement of cryo-EM and cryo-ET maps by local quality-aware deep learning with Mamba

**DOI:** 10.1101/2025.09.03.674102

**Authors:** Hong Cao, Yueting Zhu, Tao Li, Ji Chen, Jiahua He, Xinggang Wang, Sheng-You Huang

**Author notes:** (S.H.), (X.W.), (J.H.). These authors contribute equally to the work (Hong Cao, Yueting Zhu).

## Abstract

Cryo-electron microscopy (cryo-EM) has emerged as a leading technology for determining the structures of biological macromolecules. However, map quality issues such as noise and loss of contrast hinder accurate map interpretation. Traditional and deep learning-based post-processing methods offer improvements but face limitations particularly in handling map heterogeneity. Here, we present a generalist Mamba-based deep learning model for improving cryo-EM maps, named EMReady2. EMReady2 introduces a fast Mamba-based dual-branch UNet architecture to jointly capture local and global features. In addition, EMReady2 also uses a local resolution-guided learning strategy to address map heterogeneity, and significantly extends the training set. These advances render EMReady2 applicable to a broader range of cryo-EM maps, including those containing nucleic acids, medium-resolution maps, and cryo-electron tomography (cryo-ET) maps, while substantially reducing computational cost. EMReady2 is extensively evaluated on 136 diverse maps at 2.0–10.0 Å resolutions, and compared with existing map post-processing methods. It is shown that EMReady2 exhibits state-of-the-art performance in both map quality and map interpretability improvement. EMReady2 is freely available at https://github.com/huang-laboratory/EMReady2/.

## 1 Introduction

The rapid development of hardware^1^, sample preparation^2^ and image processing algorithms^3–6^ has positioned cryo-EM as one of the mainstream technologies for elucidating structural basis of biological molecules as well as the interactions among them^7–9^. In order to build accurate atomic models of biological macromolecules using cryo-EM^10–20^, the quality and interpretability of the reconstructed three-dimensional (3D) density maps is critical. However, owing to radiation damage, particle flexibility, preferred orientation, and reconstruction error, the original reconstructed 3D density maps often suffer from a high level of background noise, loss of contrast at high frequencies, as well as heterogeneity of resolution across the map^21–24^, which pose significant obstacles for accurate structural interpretation.

To reduce these defects and improve the quality of cryo-EM density maps, various post-processing algorithms have been developed, which can be broadly categorized into two classes: traditional methods^25–30^ and deep learning-based methods^31–34^. Traditional approaches for map post-processing are often performed in the Fourier space of density maps, where Fourier coefficients/amplitudes at different frequencies are re-weighted to enhance contrast and reduce noise. Leveraging the advancements in deep learning algorithms and their proven efficacy in image restoration tasks, deep learning-based post-processing methodologies have also been developed recently^31–34^. Unlike traditional map sharpening methods, deep learning-based post-processing directly modifies the input map in real space, offering the advantage of leveraging complex, data-driven models to accurately capture and enhance underlying structural details while maintaining the ability of contrast enhancement and noise reduction.

Among current deep learning approaches, EMReady exhibits the state-of-the-art performance^33^, and has been widely used to improve cryo-EM density maps for accurate map interpretations^34–38^. However, EMReady still faces limitations. First, one major limitation is that EMReady neglects the inherent density heterogeneity in cryo-EM maps during training as it generates the target simulated map by applying a uniform resolution across the entire structure. Second, EMReady mainly focuses on cryo-EM maps for proteins, and therefore its performance is limited for nucleic acid cases. Third, EMReady is trained with maps of 3–6 Å resolutions and therefore not effective enough for lower-resolution maps like cryo-ET subtomogram averaging (STA) maps. Finally, the Swin-Conv-UNet (SCUNet) architecture of EMReady is relatively slow in computation and high demand for GPU memory, which introduces a heavy computational burden for large maps.

Addressing the limitations, we propose EMReady2, a heterogeneity-aware deep learning-based post-processing method for protein and nucleic acids maps at 2–10 Å resolutions by taking advantage of the latest Bidirectional Mamba network architecture. Compared with the previous EMReady method, several major improvements have been made in EMReady2. First, the density heterogeneity is considered by constructing local resolution-dependent simulated maps during training. Second, protein and nucleic acid maps are well balanced in the training set. Third, lower-resolution (i.e. 6–10Å) maps including cryo-ET cases are added in the training set. Finally, a novel network, bidirectional Mamba-conv UNet (BiMCUnet), which integrates a bidirectional Mamba branch ^44, 45^ and a convolutional branch within a UNet architecture, is developed to greatly accelerate the computational efficiency without sacrificing the accuracy. EMReady2 is extensively validated on 136 cryo-EM and cryo-ET maps of proteins and nucleic acids with resolutions ranging from 2.0 to 10.0 Å. The evaluation results demonstrate that EMReady2 achieves significant improvements compared to existing methods including EMReady.

## 2 Results

### 2.1 Overview of EMReady2

Figure 1 shows an overview of EMReady2. The input for EMReady2 is a single EM density map, which can originate from either cryo-EM single-particle analysis (SPA) or cryo-electron tomography subtomogram averaging (STA) and may include proteins, nucleic acids, or their complexes. The input map is first partitioned into a series of overlapping volumetric slices. These slices are then fed into the BiMCUNet network to generate corresponding post-processed slices. Finally, the processed slices are reassembled to produce the output map.

**Figure 1.**
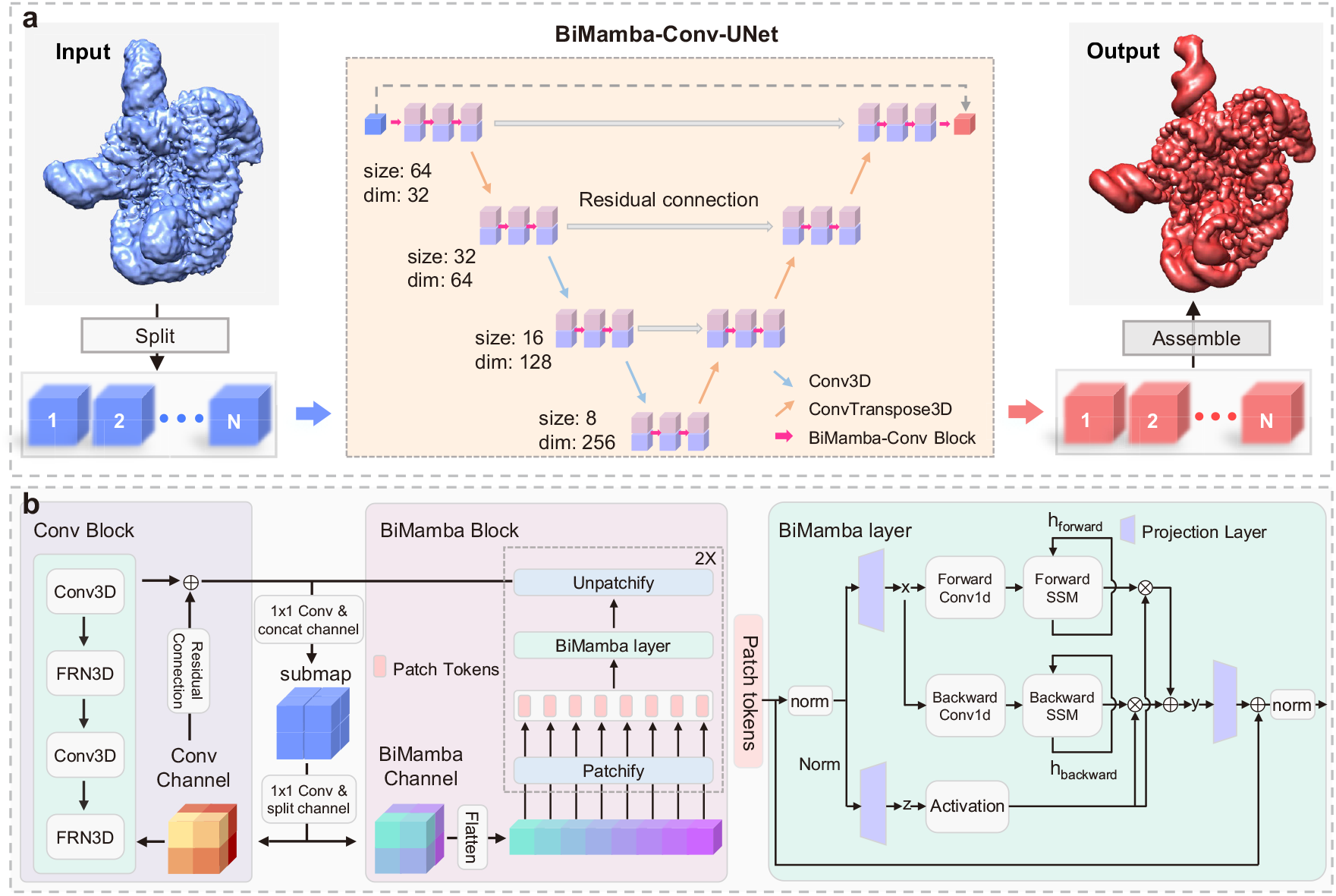
The overview of EMReady2. **a**, The workflow of EMReady2. The input to EMReady2 is an electron microscopy (EM) density map derived from cryo-EM single-particle analysis (SPA) or cryo-ET subtomogram averaging (STA), which may include proteins, nucleic acids, or their complexes. Initially, the density map is partitioned into a series of overlapping volumetric slices. These slices are then processed using the BiMamba-Conv-UNet model to generate enhanced, post-processed slices. Finally, the processed slices are reassembled to produce the refined output density map. **b**, BiMamba-Conv block architecture. The input features are processed through both the bidirectional Mamba block and the convolutional block, followed by feature fusion. The input features to the bidirectional Mamba block are first partitioned into a series of smaller patches and then projected into patch tokens. Unlike the original Mamba, the BiMamba layer processes the token sequence in both forward and backward directions, enabling more comprehensive contextual learning.

Given that convolutional layers excel at extracting local features while transformers capture global context but suffer from quadratic complexity^44, 45^, we were inspired by the remarkable success of Mamba in language modeling to introduce it–innovatively and for the first time–into the cryo-EM processing pipeline. However, the original Mamba design, tailored for sequential data, presents two key challenges in this context: unidirectional modeling and limited sensitivity to local structural details. To address these challenges, we developed the BiMCUNet, which employs bidirectional state space models (SSMs)^40–45^ to achieve comprehensive global context modeling while seamlessly integrating convolutional modules to enhance local feature extraction. Figure 1a shows the overall network architecture, consisting of three encoders, a bottleneck, and three decoders connected via skip connections. Figure 1b details the workflow within each bidirectional Mamba-conv (BiMC) Block. Further methodological details are provided in the Methods section.

### 2.2 Overall performances

We first evaluated the performance of EMReady2 on a test set of 118 cryo-EM maps at 2–10 Å. As shown in Fig. 2a, EMReady2 markedly improves the unmasked map-model FSC-0.5 relative to the deposited maps, outperforming all other post-processing methods. It should be noticed that by default we calculate the unmasked FSC in this work, unless otherwise specified. Specifically, EMReady2 improves the FSC-0.5^27^ for 113 out of the 118 maps in the test set. On average, EMReady2 achieves an FSC-0.5 of 4.65 Å, compared with 5.78 Å for the deposited maps, 5.41 Å for DeepEMhancer, 5.78 Å for phenix.auto_sharpen, 5.04 Å for EMReady, and 5.08 Å for CryoTEN (Table 1). In addition to FSC-0.5, the Q-score^46^ of maps processed by EMReady2 is also substantially improved (Fig. 2b). Specifically, the mean Q-score for EMReady2 is 0.493, compared with 0.454 for the deposited maps, 0.368 for DeepEMhancer, 0.449 for phenix.auto_sharpen, 0.477 for EMReady, and 0.452 for CryoTEN (Table 1).

**Table 1.**
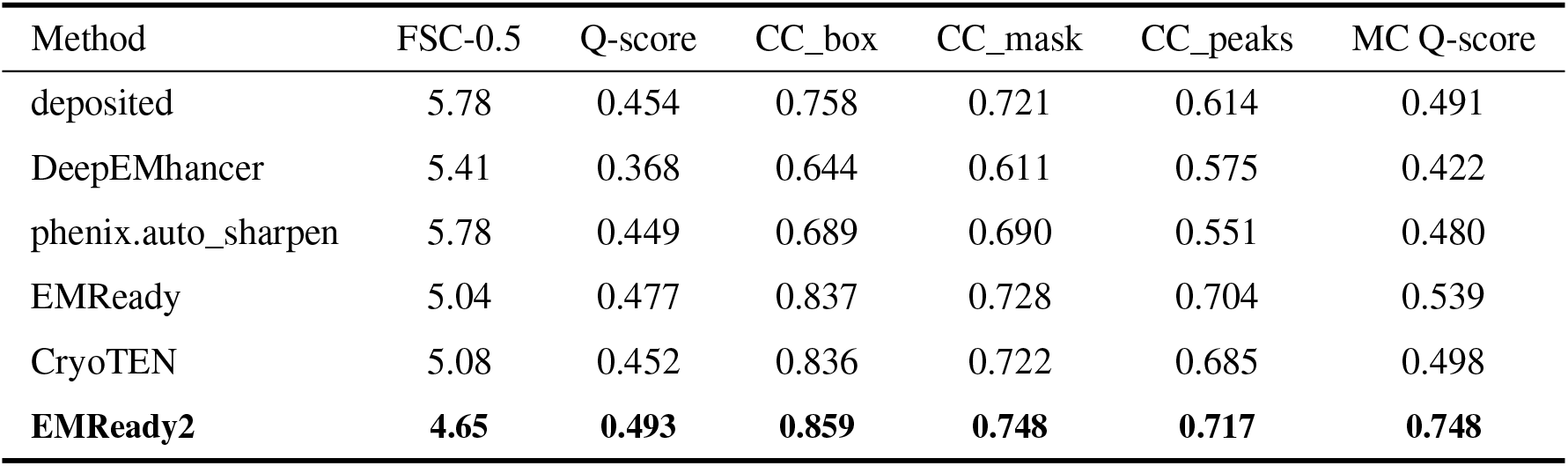
Comparison of the qualities for the deposited and processed maps on a test set of 118 cryo-EM maps at 2–10 Å resolutions in terms of FSC-0.5, Q-score, CC_box, CC_mask, CC_peaks, and main-chain (MC) Q-score. Here, the lower value is the better for FSC-0.5, while the higher value is the better for the other metrics.

| Method | FSC-0.5 | Q-score | CC_box | CC_mask | CC_peaks | MC Q-score |
| --- | --- | --- | --- | --- | --- | --- |
| deposited | 5.78 | 0.454 | 0.758 | 0.721 | 0.614 | 0.491 |
| DeepEMhancer | 5.41 | 0.368 | 0.644 | 0.611 | 0.575 | 0.422 |
| phenix.auto_sharpen | 5.78 | 0.449 | 0.689 | 0.690 | 0.551 | 0.480 |
| EMReady | 5.04 | 0.477 | 0.837 | 0.728 | 0.704 | 0.539 |
| CryoTEN | 5.08 | 0.452 | 0.836 | 0.722 | 0.685 | 0.498 |
| **EMReady2** | **4.65** | **0.493** | **0.859** | **0.748** | **0.717** | **0.748** |

**Figure 2.**
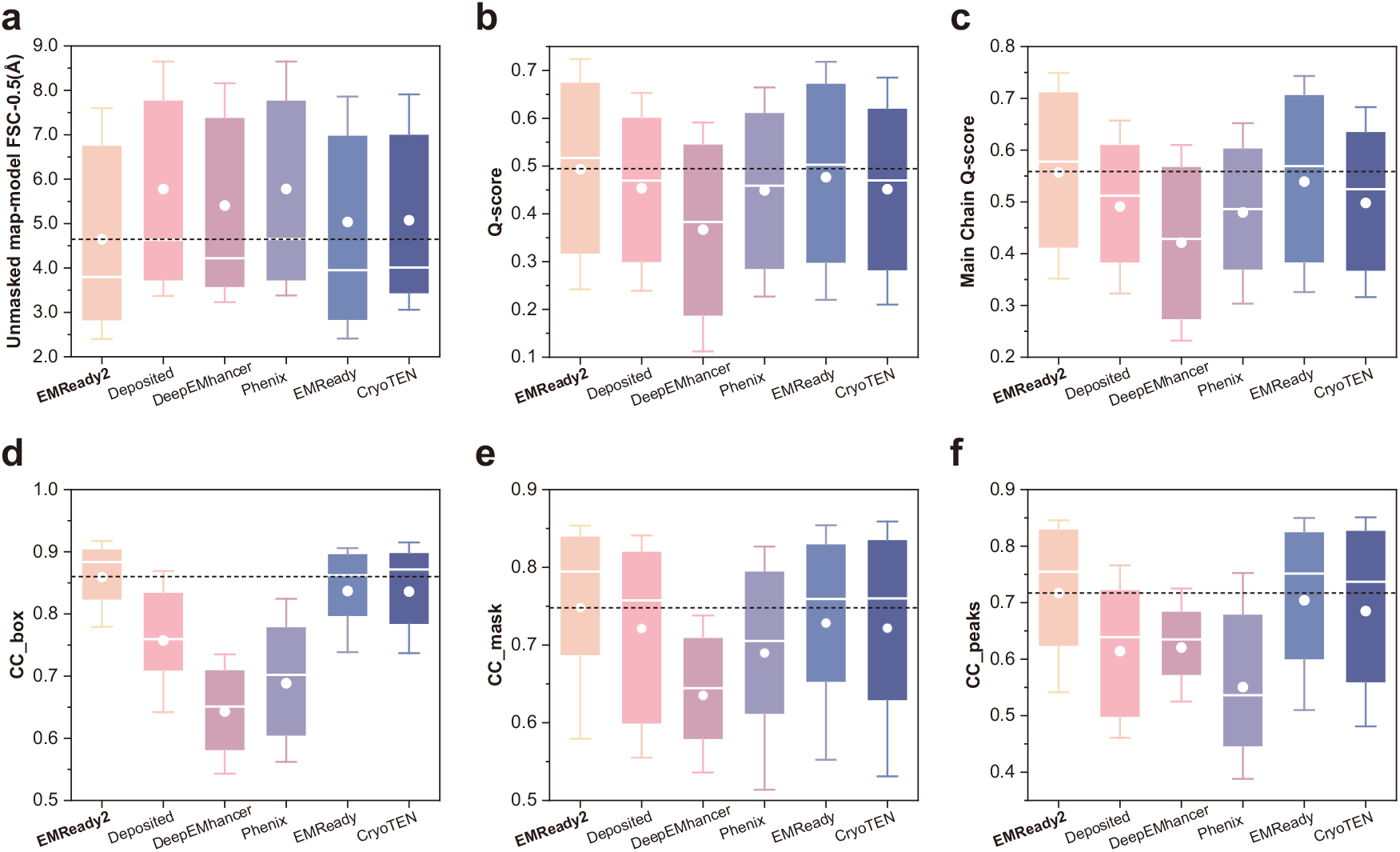
Evaluation results on 118 primary cryo-EM maps. **a-f**, Box-and-whisker plots of unmasked map–model FSC-0.5 (**a**), Q-score (**b**), main chain Q-score (**c**), CC_box (**d**), CC_mask (**e**), CC_peaks (**f**) for the EMReady2-processed, deposited, DeepEMhancer-processed, phenix.auto_sharpen-processed, EMReady-processed, CryoTEN-processed maps. The central line represents the median, with circles indicating the mean value. The lower and upper hinges correspond to the first and third quartiles, respectively, while the whiskers extend to 1.5 times the interquartile range. The dashed line indicates the mean value of the EMReady2-processed map. Here, the lower value is the better for FSC-0.5, while the higher value is the better for the other metrics.

In addition, we further assessed the main chain (MC) Q-score to characterize map continuity (Fig. 2c), which is critical for reliable backbone tracing in subsequent model building. It is shown that EMReady2 improves the main chain Q-score in 115 of the 118 maps. On average, EMReady2 achieves a main chain Q-score of 0.748, which is considerably higher than 0.491 of the deposited map (Table 1). Even when compared with EMReady, which yields the best performance among other methods (with a main chain Q-score of 0.539), EMReady2 still achieves an improvement of nearly 20 percentage points. Furthermore, EMReady2 is also capable of improving CC_box, CC_mask, and CC_peaks. The maps processed by EMReady2 achieve average CC_box, CC_mask, and CC_peaks values of 0.859, 0.748, and 0.717, respectively, which are significantly higher than 0.758, 0.721, and 0.614 for the deposited map (Table 1), and those of the other post-processing approaches including EMReady.(Fig. 2d-f). Specifically, EMReady2 improved CC_box in 109 maps, CC_mask in 92 maps, and CC_peaks in 101 maps within the test set.

Besides single particle analysis (SPA) approach, another important domain of EM-based structural biology is cryo-electron tomography (cryo-ET), which produce density maps at sub-nanometre resolution through subtomogram averaging (STA). We further evaluated the performance of EM-Ready2 on a test set of 18 cryo-ET STA maps at 3–10 Å, and compared it with DeepEMhancer, phenix.auto_sharpen, EMReady, and CryoTEN. Given that most cryo-ET STA maps in this set contain only partially modeled regions, we selected masked map-model FSC-0.5, Q-score, CC_mask, and main chain Q-score as evaluation metrics. As shown in Table 2, EMReady achieves a masked map-model FSC-0.5 of 6.26 Å, a Q-score of 0.396, a CC_box of 0.698, and a main-chain Q-score of 0.451, all substantially better than those for the deposited map (i.e. 7.06 Å, 0.360, 0.673, and 0.397, respectively) as well as other methods. These results demonstrates the robust performance of EMReady2 on STA data.

**Table 2.** Comparison of the qualities for the deposited and processed maps on the test set of 18 cryo-ET maps at 3-10 Å resolutions in terms of FSC-0.5, Q-score, CC_box, CC_mask, CC_peaks, and main-chain (MC) Q-score. Here, the lower value is the better for FSC-0.5, while the higher value is the better for the other metrics.

| Method | Masked FSC-0.5 | Q-score | CC_box | MC Q-score |
| --- | --- | --- | --- | --- |
| deposited | 7.06 | 0.360 | 0.673 | 0.397 |
| DeepEMhancer | 6.81 | 0.305 | 0.645 | 0.359 |
| phenix.auto_sharpen | 6.92 | 0.364 | 0.629 | 0.399 |
| EMReady | 6.45 | 0.379 | 0.654 | 0.428 |
| CryoTEN | 6.91 | 0.348 | 0.624 | 0.395 |
| **EMReady2** | **6.26** | **0.396** | **0.698** | **0.451** |

Figure 3 shows a representative example using EMD-6551, a 3.8 Å cryo-EM map of the magnesium channel CorA in its closed, symmetric magnesium-bound state. We conducted a detailed analysis across different contour levels to illustrate the improvements introduced by EMReady2. At low contour levels, EMReady2 effectively enhances the contrast between the macromolecule and the surrounding lipid nanodisc, rendering the unmodeled lipid regions nearly invisible in the processed map. This highlights both the denoising and global optimization capabilities of EMReady2, which are also reflected in the markedly improved Fourier shell correlation (FSC) shown in Fig. 3b. Quantitatively, the FSC-0.5 of the EMReady2-processed map reaches 3.56 Å, a substantial improvement over the 4.57Å for the deposited map. We then performed a comparison at the medium contour level. As shown in Fig. 3c, the enlarged view in the middle displays the details of the transmembrane region of the ion channel, from which the improvement in side-chain density by EMReady2 is distinctly observable. This is reflected by the Q-score of the EMReady2-processed map, which is 0.646, higher than 0.550 for the deposited map. At high contour levels, the zoomed-in panel highlights the alignment between the backbone density and the corresponding atomic model (PDB: 3JCF). EM-Ready2 notably enhances the continuity and clarity of the backbone trace, which is consistent with its substantial improvement in the main chain Q-score from 0.584 for the deposited map to 0.786 for the EMReady2-processed map. In addition, EMReady2 yields higher CC metrics, with CC_box, CC_mask, and CC_peaks reaching 0.906, 0.826, and 0.832, respectively, substantially outperforming the deposited map values of 0.742, 0.814, and 0.660.

**Figure 3.**
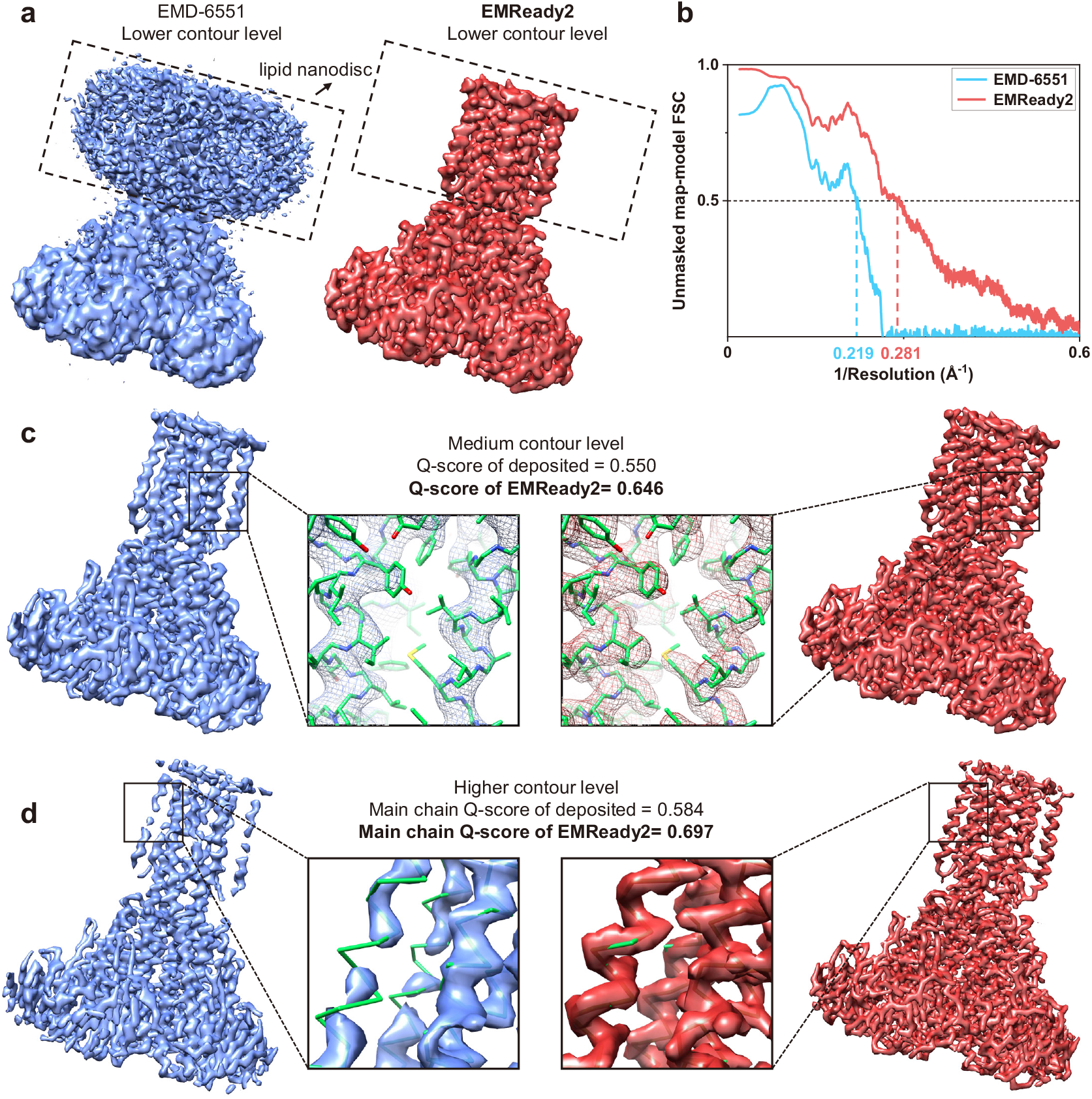
The EMReady2 results on map EMD-6551 (PDB ID: 3JCF) at 3.80 Å resolution. The deposited primary maps are shown in blue, the EMReady2-processed maps in red, and the PDB structures in green. **a**, Comparison of the deposited and EMReady2-processed maps at a low contour level. Boxes highlight regions of unmodeled lipids. **b**, Unmasked map-model Fourier shell correlation as a function of inverse resolution for both the deposited and EMReady2-processed maps. **c**, Comparison of the deposited and EMReady2-processed maps at a medium contour level. The central insets highlight the ion channel region, with particular focus on the side chain structures and their fit to the density map. **d**, Comparison of the deposited and EMReady2-processed maps at a high contour level. The central insets highlight the main chain density continuity. Contours are drawn to enclose equal volumes for each case.

### 2.3 Improvement of nucleic acid maps

To evaluate the ability of EMReady2 in improving nucleic acid density of electron microscopy (EM) maps, we further assessed its performance on a test set of 18 raw cryo-EM maps in which nucleic acid residues account for more than 10% of the corresponding structures. EMReady2 was compared against DeepEMhancer, phenix.auto_sharpen, EMReady, and CryoTEN. Performance was quantified using six distinct metrics: FSC-0.5, CC_box, CC_mask, CC_peaks, Q-score, and main chain Q-score.

As shown in Fig. 4a, EMReady2 achieves a mean Q-score of 0.499, which is significantly higher than 0.453 for the deposited maps, 0.368 for DeepEMhancer, 0.446 for phenix.auto_sharpen, 0.458 for EMReady and 0.445 for CryoTEN (Table 3). Figure 4b,c show the results of FSC-0.5 across different methods. Overall, EMReady2 obtains a mean value of 4.20 Å, representing a substantial improvement over the 5.70 Å for deposited maps, 4.94 Å for DeepEMhancer, 5.70 Å for phenix.auto_sharpen, 4.59 Å for EMReady, and 4.69 Å for CryoTEN (Table 3). Examining individual cases in the test set further demonstrates that EMReady2 exhibits consistent and broadly applicable improvements. Figure 4d,e,f shows the average map-model correlation coefficients (CC_box, CC_mask, and CC_peaks). Specifically, the average CC_box, CC_mask, and CC_peaks values for the maps processed by EM-Ready2 are 0.889, 0.786, and 0.767, respectively, which are substantially higher than 0.820, 0.765, and 0.706 for the deposited maps (Table 3). Considering all the evaluation metrics, EMReady and EMReady2 are both capable of improving the deposited primary maps, and significantly outperform DeepEMhancer and phenix.auto_sharpen. Furthermore, EMReady2 achieves a superior performance to EMReady, suggesting the effectiveness of including local quality information and nucleic acid data during training. In terms of main chain Q-score, EMReady2 also shows a striking performance, achieving a value of 0.686, an improvement of over 18 percentage points compared with 0.501 for the best result among the other methods, EMReady.

**Table 3.** Comparison of the qualities for the deposited and processed maps on a test set of 18 cryo-EM maps with nucleic acid content exceeding 10% in terms of FSC-0.5, Q-score, CC_box, CC_mask, CC_peaks, and main-chain (MC) Q-score. Here, the lower value is the better for FSC-0.5, while the higher value is the better for the other metrics.

| Method | FSC-0.5 | Q-score | CC_box | CC_mask | CC_peaks | MC Q-score |
| --- | --- | --- | --- | --- | --- | --- |
| deposited | 5.70 | 0.453 | 0.820 | 0.765 | 0.706 | 0.462 |
| DeepEMhancer | 4.94 | 0.368 | 0.665 | 0.659 | 0.650 | 0.400 |
| phenix.auto_sharpen | 5.70 | 0.446 | 0.702 | 0.704 | 0.582 | 0.448 |
| EMReady | 4.59 | 0.458 | 0.871 | 0.771 | 0.761 | 0.501 |
| CryoTEN | 4.69 | 0.445 | 0.864 | 0.752 | 0.735 | 0.470 |
| **EMReady2** | **4.20** | **0.499** | **0.889** | **0.786** | **0.767** | **0.686** |

**Figure 4.**
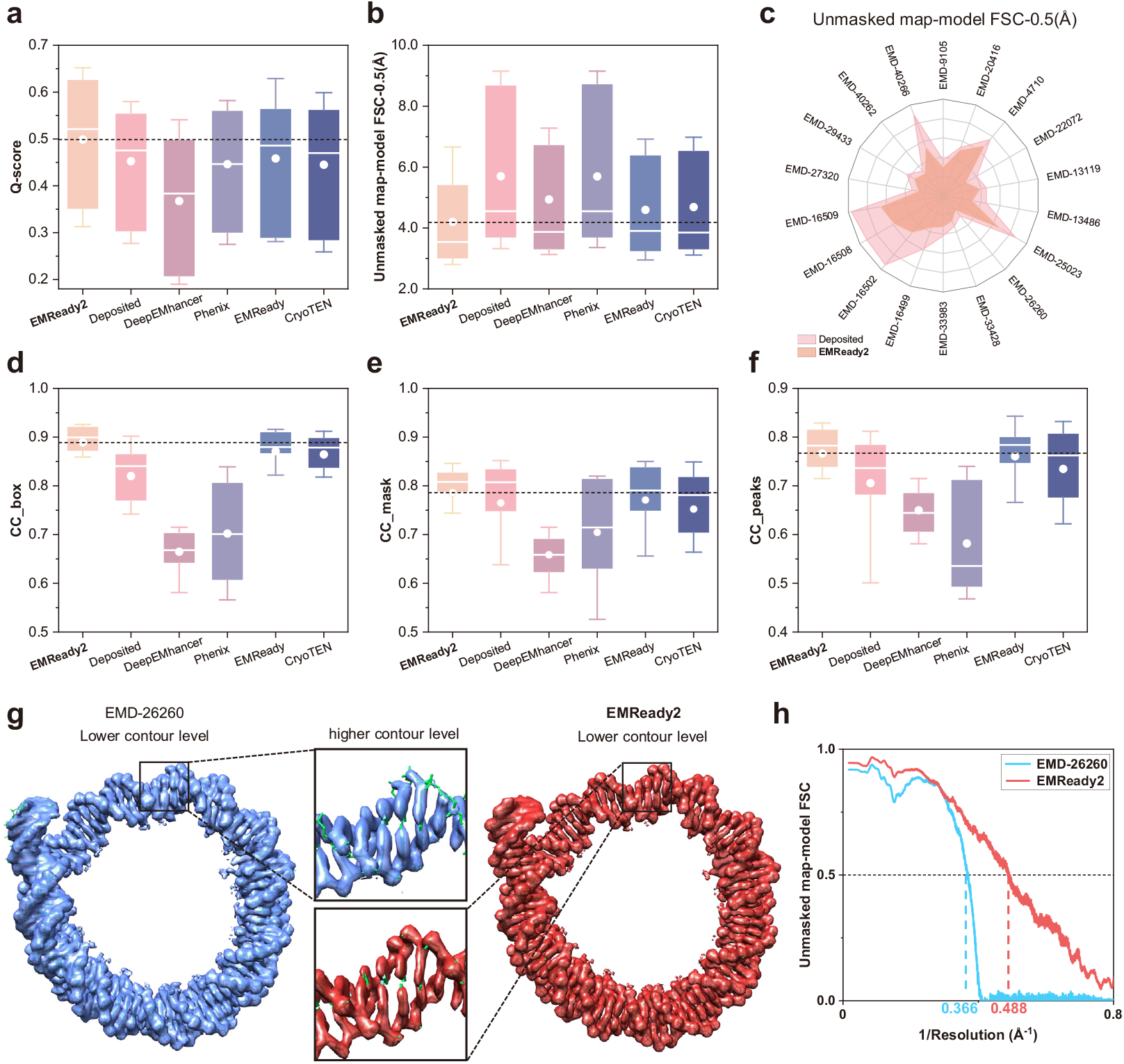
Evaluation results on the test set of 18 cryo-EM maps with nucleic acids exceeding 10%. **a,b,d-f**, Box-and-whisker plots showing unmasked map–model Q-score (a), FSC-0.5 (**b**), CC_box (**d**), CC_mask (**e**), and CC_peaks (**f**) for EMReady2-processed, deposited, DeepEMhancer-processed, phenix.auto_sharpen-processed, EMReady-processed, and CryoTEN-processed maps. The central line represents the median, with circles indicating the mean value. The lower and upper hinges correspond to the first and third quartiles, respectively, while the whiskers extend to 1.5 times the interquartile range. The dashed line represents the mean value of the EMReady2-processed map. **c**, Comparison of the unmasked map-model FSC-0.5 between the deposited map and the EMReady2-processed map for each test case. **g**, Example of EMD-26260 (PDB ID: 7U0I) at 2.6 Å resolution. The deposited primary maps are shown in blue, the EMReady2-processed maps in red, and the PDB structures in green. Contours are drawn to enclose equal volumes for each case. **h**, Unmasked map-model Fourier shell correlation as a function of inverse resolution for the deposited maps and the EMReady2-processed map on EMD-26260.

Figure 4g,h show an example of EMD-26260, a 2.6 Å cryo-EM map of the LIN28b nucleosome bound to OCT4. At low contour level, EMReady2 effectively preserves the high-quality regions of the original map while refining side-chain densities. At higher contour levels, EMReady2 also demonstrates impressive improvements in the nucleic acid backbone density. The FSC-0.5 curves further confirm the enhancements across all spatial frequencies in Fourier space. Quantitatively, the EMReady2-processed map achieves an improved Q-score of 0.740, significantly higher than 0.698 for deposited map, 0.656 for DeepEMhancer, 0.701 for phenix.auto_sharpen, 0.720 for EMReady, and 0.710 for CryoTEN. EMReady2 also achieves the highest unmasked map–model FSC-0.5 value of 2.05 Å.

### 2.4 Improvement in map interpretability

The true enhancement of density map quality should improve the accuracy of the built models, or namely enhance the interpretability of the maps. Therefore, we conducted further analysis to assess the impact of EMReady2 in model building. To ensure unbiased results, we utilized phenix.map_to_model^27^ for automatic building of atomic models from the map regions segmented within 4.0 Å. To ensure reliable execution of phenix.map_to_model, we selected only those maps with resolutions better than 5.0 Å, yielding a total of 832 chains comprising 753 protein chains and 79 nucleic acid chains. The built models are compared with the deposited PDB structures through phenix.chain_comparison^27^, from which two metrics are reported: residue coverage and sequence recall.

The evaluation results on the protein-chain test set are presented in Fig. 5a,b and summarized in Table 4. Notably, EMReady2 substantially improves the residue coverage of the built atomic models, achieving an average residue coverage of 70.51%, compared with 56.89% for the deposited maps, 53.91% for DeepEMhancer, 54.26% for phenix.auto_sharpen, 65.33% for EMReady, and 61.49% for CryoTEN (Table 4). In terms of sequence recall, EMReady2 reaches 32.24% on proteins, substantially outperforming the deposited maps at 18.31% as well as other methods. Figure 5c shows the modeling results for chain E of 6R25. Comparison of the loop regions indicates substantial improvements made by EMReady2 that lead to improved model building. On the EMReady2-processed map, the built model achieves a residue coverage and a sequence recall of 91.10% and 71.73%, respectively, which are much better than 72.70% and 3.05% for the model built on the deposited map.

**Table 4.** Comparison of the map interpretability in terms of de novo model building on the test set of 96 cryo-EM maps processed by EMReady2 and four other methods. The test set include 753 protein chains and 79 nucleic acid chains. Method Residue coverage (%) Sequence recall (%)

| Method | Residue coverage (%) |  | Sequence recall (%) |  |
| --- | --- | --- | --- | --- |
|  | Protein | Nucleic acid | Protein | Nucleic acid |
| deposited | 56.89 | 54.31 | 18.31 | 20.99 |
| DeepEMhancer | 53.91 | 50.02 | 16.67 | 18.54 |
| phenix.auto_sharpen | 54.26 | 57.14 | 18.03 | 20.77 |
| EMReady | 65.33 | 61.56 | 30.00 | 24.87 |
| CryoTEN | 61.49 | 59.01 | 23.27 | 22.97 |
| **EMReady2** | **70.51** | **72.14** | **32.24** | **27.40** |

**Figure 5.**
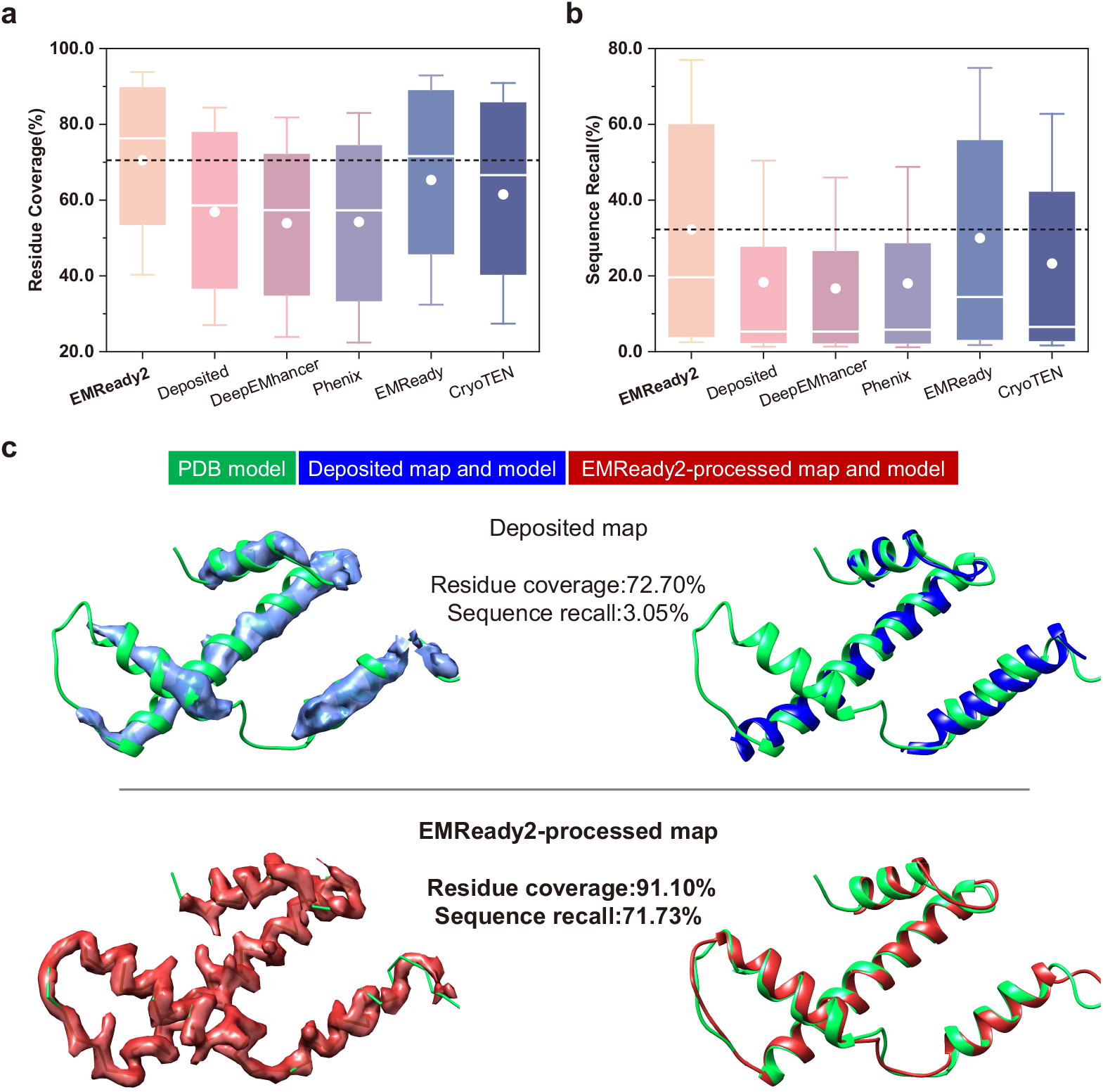
Improvement in map interpretability on the test set of 753 protein chains. **a,b**, Box- and-whisker plots showing residue coverage (**a**) and sequence recall (**b**) for the built models of 753 protein chains. The central line represents the median, with circles indicating the mean value. The lower and upper hinges correspond to the first and third quartiles, respectively, while the whiskers extend to 1.5 times the interquartile range. The dashed line represents the mean value of the EMReady2processed map. **c**, Chain E of 6R25 on EMD-4710 at 4.61 Å resolution. The upper and lower panels display the deposited and the EMReady2-processed maps and their corresponding built models, respectively. The reference PDB structures are colored in green. The deposited maps and their corresponding models are colored in blue. The EMReady2-processed maps and their corresponding models are colored in red. Contours are drawn to enclose equal volumes for each case.

The evaluation results on the nucleic-acid-chain test set are shown in Fig. 6a,b and summarized in Table 4. For nucleic acids, EMReady2 achieves even more improvements in residue coverage, with an average of 72.14%, compared with 54.31% for the deposited maps, 50.02% for DeepEMhancer, 57.14% for phenix.auto_sharpen, 61.56% for EMReady, and 59.01% for CryoTEN. Although the sequence types in nucleic acids are generally more challenging to predict, EMReady2 still yields an improvement in sequence recall, reaching an average of 27.40%, higher than 20.99% for the deposited maps and other methods. Figure 6c compares the modeling results for chains I and J of 6R25 (EMD-4710). It can be seen from the figure that EMReady2 effectively enhances the density traces for the DNA double helix, and thus results in a better-built model. Specifically, the model built on the EMReady2-processed map achieves a residue coverage of 81.60% and a sequence recall of 24.85%, respectively, compared with only 19.70% and 5.77% for the deposited maps.

**Figure 6.**
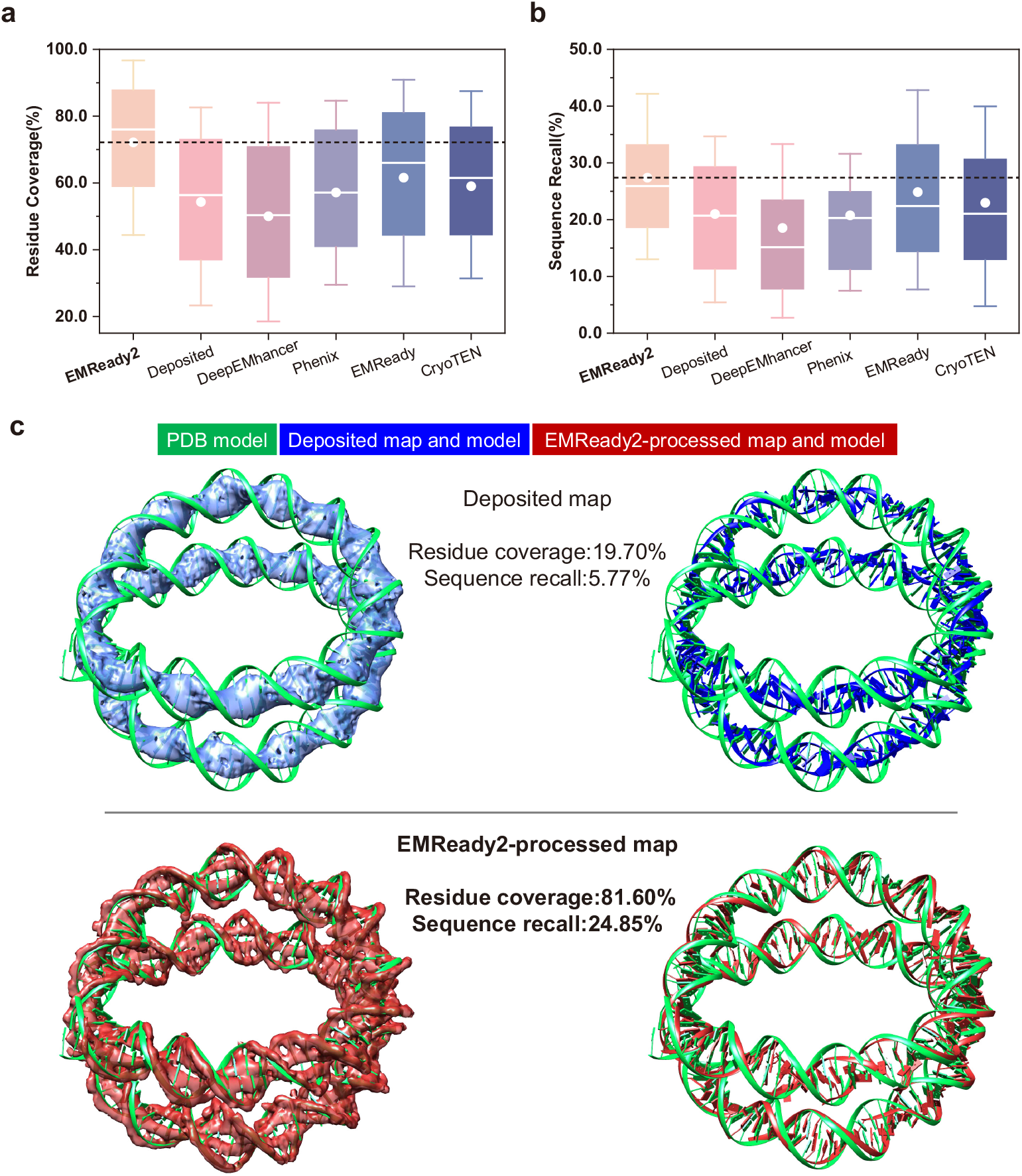
Improvement in map interpretability on the test set of 79 nucleic acid chains. **a,b**, Box-and-whisker plots showing residue coverage (**a**) and sequence recall (**b**) for the built models of 79 nucleic acid chains. The central line represents the median, with circles indicating the mean value. The lower and upper hinges correspond to the first and third quartiles, respectively, while the whiskers extend to 1.5 times the interquartile range. The dashed line represents the mean value of the EMReady2-processed map. **c**, Chain I and J of 6R25 on EMD-4710 at 4.61 Å resolution. The upper and lower panels display the deposited and the EMReady2-processed maps and their corresponding built models, respectively. The reference PDB structures are colored in green. The deposited maps and their corresponding models are colored in blue. The EMReady2-processed maps and their corresponding models are colored in red. Contours are drawn to enclose equal volumes for each case.

### 2.5 Robustness against map heterogeneity and anisotropy

Intrinsic flexibility of the target molecule, preferred orientation of the particles on the grid, or errors in the subsequent image analysis, can lead to heterogeneity in the reconstructed EM density maps^21–23^. Here, we carry out detailed analysis of how EMReady2 improves the maps with heterogeneity.

Figure 7a,b shows the example of EMD-33428, a 3.01 Å cryo-EM map of Native Tetrahymena ribozyme. Comparison of the local resolution maps calculated using MonoRes^21^ for the deposited map and the EMReady2-processed map reveals pronounced signal heterogeneity between the protein and nucleic acid regions in the deposited map (the left panel of Fig. 7a). After post-processing, EM-Ready2 not only effectively enhances the high-resolution protein densities but also recovers certain low-resolution features of the nucleic acids, thereby mitigating the overall heterogeneity of the density map to some extent (the right panel of Fig. 7a). The FSC curve displayed in Fig. 7b also demonstrates that EMReady2 is able to enhance the signals along a broad range of spatial frequencies.

**Figure 7.**
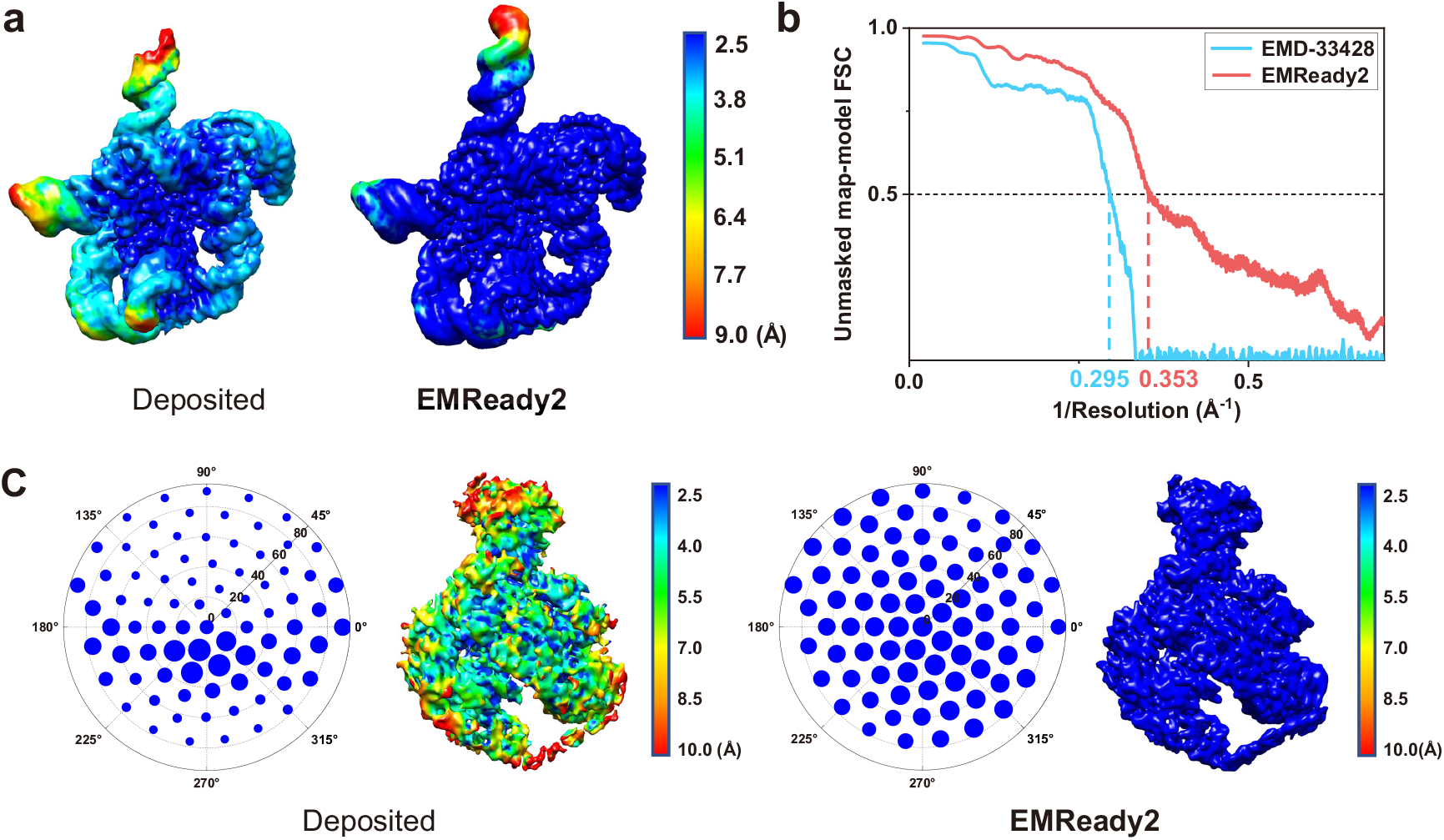
Robustness of EMReady2 against map heterogeneity and anisotropy. **a**, Example of EMD-33428 (associated PDB ID: 7XSN) at 3.01 Å resolution. Comparison of local resolution maps between the deposited map (left panel) and the EMReady2-processed map (right panel). **b**, Unmasked map–model Fourier shell correlation as a function of inverse resolution for EMD-33428 and the EMReady2-processed map. **c**, Example of EMD-20501 (PDB ID: 6PW9) at 4.03 Å resolution. The left panel shows the local resolution map and angular plots for the local-directional resolution map of the deposited map. The right panel shows the corresponding maps and plots for the EMReady2-processed map. Contours are drawn to enclose equal volumes for each case.

Figure 7c shows another example, EMD-20501, a 4.03 Å cryo-EM map of the human NatE/HYPK complex. It can be seen from the figure that EMReady2 significantly enhances the entire map including the weaker density signal at the bottom part of the map. The angular distribution of local directional resolution measured by MonoDir^22^ indicates a strong angular anisotropy in the deposited map, which is consistent with its heterogeneity of local resolution. After being processed by EMReady2, the heterogeneity in local resolution and angular resolution can be effectively mitigated.

### 2.6 Ablation experiments

Compared with EMReady, EMReady2 exhibits significant differences in three key aspects: the training framework (encompassing network architecture and input density box size), the training dataset (incorporating a mix of diverse data types), and the generation method of simulated maps (utilizing local resolution). To investigate how these factors influence the performance of EMReady, we conducted extensive ablation experiments, including: generating simulated maps using the reported resolution, adopting the same SCUnet network architecture as EMReady, excluding medium-resolution density maps from training, excluding cryo-ET density maps from training, and modifying the input box size.

We first performed ablation experiments on the test set of 118 cryo-EM maps. Five ablation models are presented in Table 5, from which it can be observed that the baseline model achieves the best performance in all metrics, although some ablation models also demonstrate competitive performance in terms of CC (correlation coefficient) values. Figure 8a,b ishow the performance comparisons of all models in terms of FSC-0.5 and Q-score, revealing the key factors affecting performance. First, the generation of simulated maps based on local resolution endows the model with enhanced capability for personalized optimization of local densities, thereby alleviating overfitting. Second, the bidirectional Mamba architecture enables efficient information interaction while maintaining high computational speed, which also allows us to train the model with a larger box size. Since the range of secondary structures can span tens of angstroms, a larger input box better captures such long-range structural information during model training, ultimately leading to improved performance. Additionally, the exclusion of either medium-resolution or cryo-ET density maps results in a significant drop in performance highlighting the necessity of training with a mixed dataset of density maps with varying resolutions. The similar phenomenon also observed in our previous work^39^. Overall, the baseline model achieves an FSC-0.5 of 4.65 Å and a Q-score of 0.493, outperforming all ablation models.

**Table 5.** The ablation experiments of EMReady2 on the test set of 118 cryo-EM maps. The five ablation models include changing the generation method of the simulated map (reported resolution), changing the network framework to SCUnet (SCUnet), not using medium-resolution density maps for training (w/o medium), not using cryo-ET density maps for training (w/o cryo-ET), and changing the input box size to 48 (box_size=48). Here, the lower value is the better for FSC-0.5, while the higher value is the better for the other metrics. The numbers in bold fonts indicate the best performances for the corresponding metrics.

| Model | FSC-0.5 | Q-score | CC_box | CC_mask | CC_peaks | MC Q-score |
| --- | --- | --- | --- | --- | --- | --- |
| **baseline** | **4.65** | **0.493** | **0.859** | **0.748** | **0.717** | **0.748** |
| reported resolution | 4.73 | 0.486 | 0.856 | 0.747 | 0.715 | 0.601 |
| SCUnet | 4.69 | 0.488 | 0.857 | 0.748 | 0.714 | 0.619 |
| w/o medium | 4.90 | 0.468 | 0.837 | 0.736 | 0.703 | 0.713 |
| w/o cryo-ET | 4.75 | 0.487 | 0.858 | 0.745 | 0.710 | 0.691 |
| box_size=48 | 4.82 | 0.478 | 0.846 | 0.740 | 0.706 | 0.590 |

**Figure 8.**
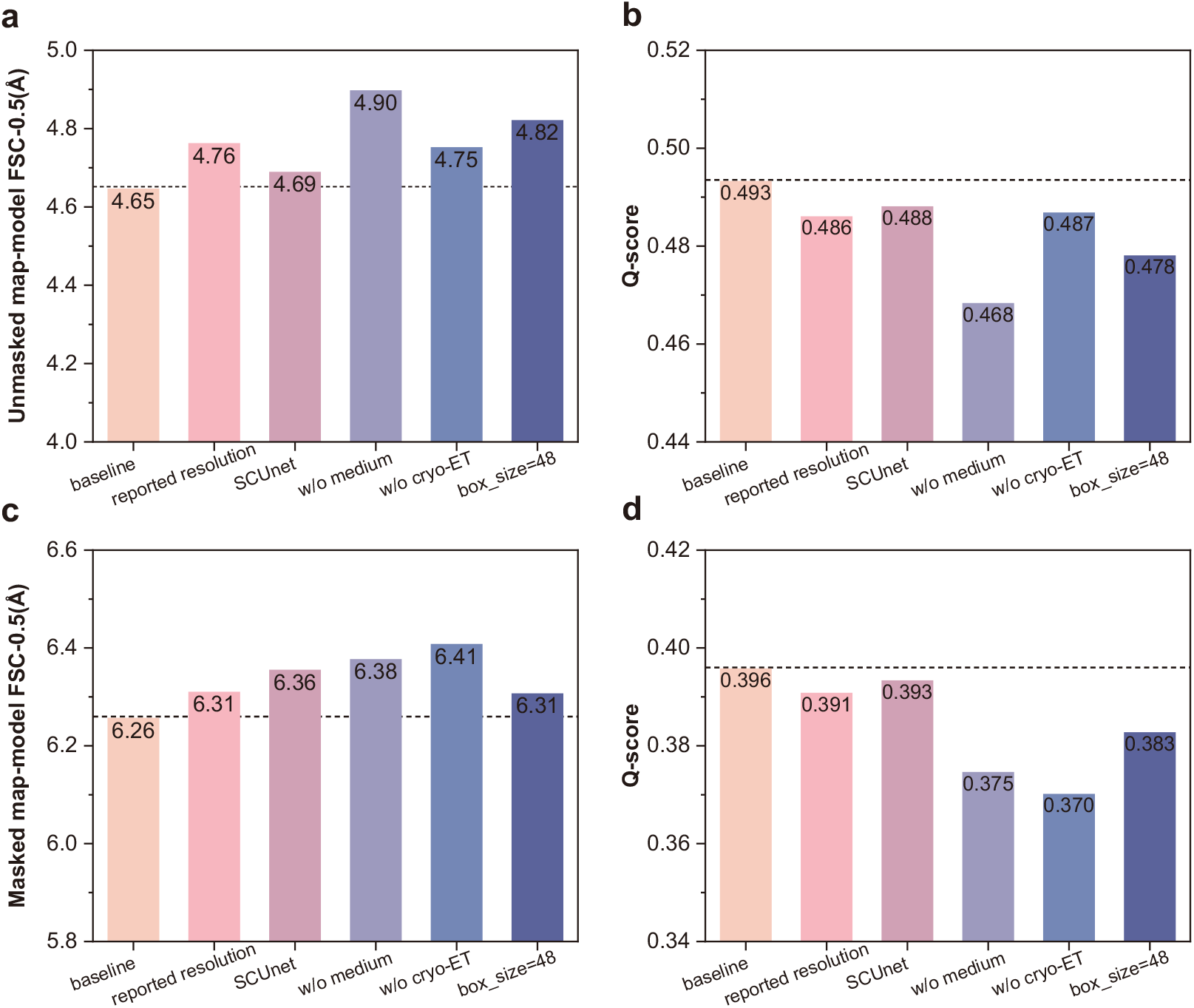
Ablation experiments of EMReady2. **a,b**, Results of the ablation experiments on the test set of 118 cryo-EM maps. The bar chart presents the ablation results for the average FSC-0.5 (**a**) and Q-score (**b**). **c,d**, Results of the ablation experiments on the test set of 18 cryo-ET maps. The bar chart presents the ablation results for the average FSC-0.5 (**c**) and Q-score (**d**). Here, the lower value is the better for FSC-0.5, while the higher value is the better for the other metrics.

Based on the aforementioned discussion, we also evaluated the model performance on a test set of 18 cryo-ET maps using four key metrics. The corresponding benchmarking results are summarized in Table 6, and the performance for each model in terms of Masked FSC-0.5 and Q-score is visualized in Fig. 8c,d. The ablation experiments on this cryo-ET test set show trends similar to those observed on the cryo-EM test set, with one notable distinction. Namely, although most cryo-ET density maps are of medium resolution, their proportion in the training dataset is relatively small (18 out of 348 maps). As such, the ‘w/o cryo-ET’ ablation model has a less pronounced impact on performance compared to the ‘w/o medium’ model in the cryo-EM test. Nevertheless, the structural patterns of cryo-ET density maps differ from those of cryo-EM density maps. Consequently, the ‘w/o cryo-ET’ ablation model exerts a more significant influence on performance in the cryo-ET test set. In summary, the baseline model has a Masked FSC-0.5 of 6.26 Å and a Q-score of 0.396, ranking as the top-performing model on this test set.

**Table 6.** The ablation experiments of EMReady2 on the test set of 18 cryo-ET maps. The five ablation models include changing the generation method of the simulated map (reported resolution), changing the network framework to SCUnet (SCUnet), not using medium-resolution density maps for training (w/o medium), not using cryo-ET density maps for training (w/o cryo-ET), and changing the input box size to 48 (box_size=48). Here, the lower value is the better for FSC-0.5, while the higher value is the better for the other metrics. The numbers in bold fonts indicate the best performances for the corresponding metrics.

| Model | Masked FSC-0.5 | Q-score | CC_mask | MC Q-score |
| --- | --- | --- | --- | --- |
| **baseline** | **6.26** | **0.396** | **0.699** | **0.451** |
| reported resolution | 6.31 | 0.391 | 0.692 | 0.410 |
| SCUnet | 6.36 | 0.393 | 0.696 | 0.425 |
| w/o medium | 6.38 | 0.375 | 0.681 | 0.395 |
| w/o cryo-ET | 6.41 | 0.370 | 0.692 | 0.409 |
| box_size=48 | 6.31 | 0.383 | 0.686 | 0.401 |

### 2.7 Computational efficiency

Figure 9 shows the running times four deep learning-based methods: EMReady2, DeepEMhancer, EMReady, and the recently introduced CryoTEN on the test set of all 136 maps. DeepEMhancer, the earliest of these methods, was released in 2021. It typically requires a substantial runtime with a mean of 90.98 s, attributable to its relatively simple yet less optimized architecture. Our previous method, EMReady, released in 2023, similarly incurs high computational costs with a mean runtime of 83.26 s, despite exhibiting strong enhancement performance. CryoTEN, introduced more recently with a focus on speed, achieves a lower average runtime of 17.33 s, though its enhancement quality remains inferior to that of EMReady. Among all tested methods, EMReady2 achieves the shortest mean runtime of 14.81 s. This represents a substantial improvement in computational efficiency over both EMReady and DeepEMhancer, while also outperforming CryoTEN in speed.

**Figure 9.**
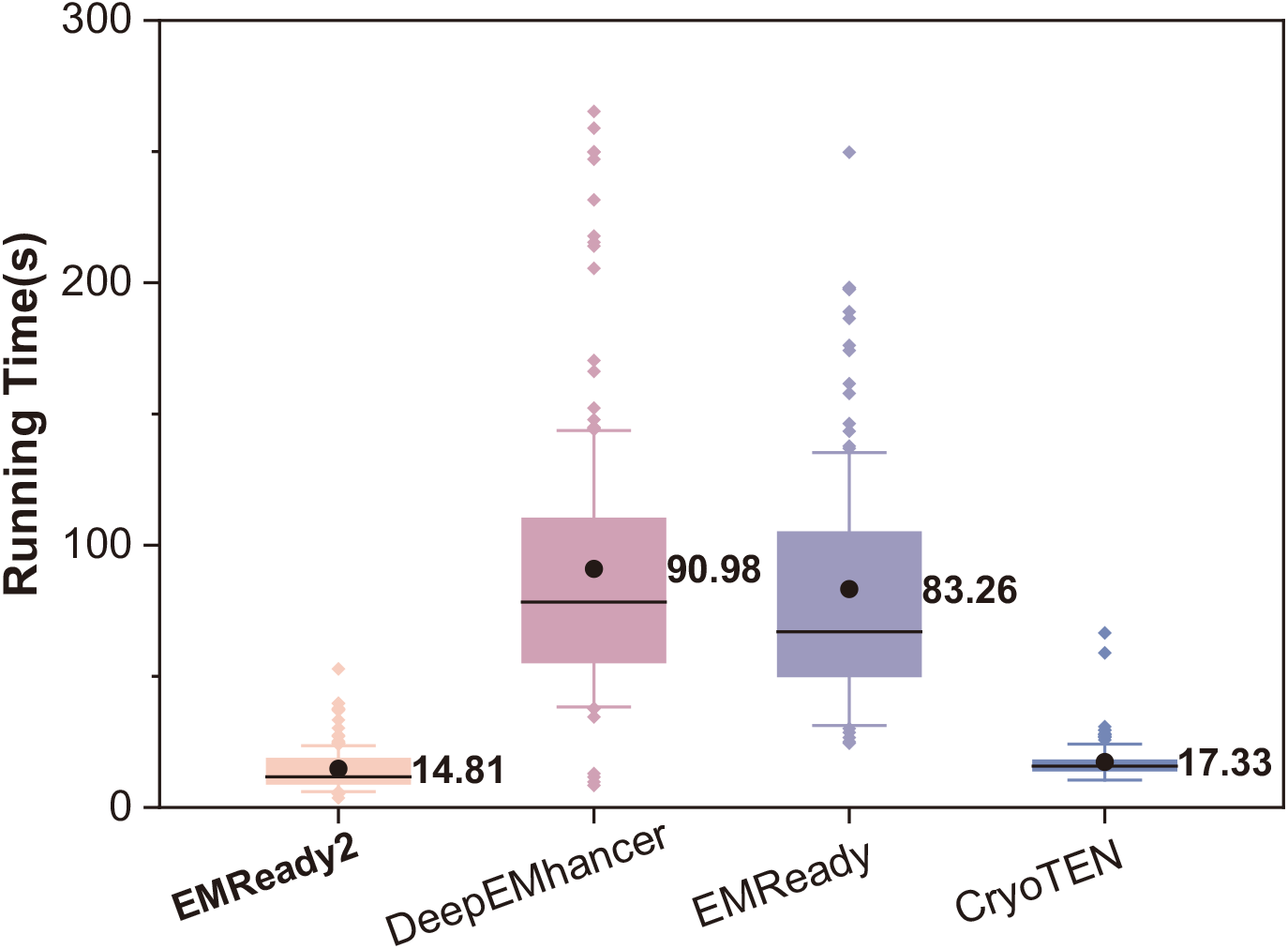
Comparison of running times for different deep learning-based methods. Box-and-whisker plots of running times on the complete test set of 136 test samples. The central line represents the median, with circles indicating the mean value. The lower and upper hinges correspond to the first and third quartiles, respectively, while the whiskers extend to 1.5 times the interquartile range. Outliers are depicted as diamonds.

## 3 Discussion

In this study, we have propose EMReady2, a universally applicable deep learning-based post-processing method for EM density maps. During training, we generate the target simulated map using varying local resolution values derived from Q-scores for different atoms within the structure. This mitigates training biases introduced by poorly modeled regions and allows the model to adapt its density modification in a heterogeneity-aware manner during inference. Furthermore, our training and evaluation now encompass maps containing nucleic acid molecules, intermediate-resolution maps, and cryo-ET maps. We also introduce Mamba for the first time into the cryo-EM processing pipeline and extend it with a bidirectional design and convolutional integration, proposing the innovative BiMamba-Conv-UNet architecture. This approach substantially enhances the extraction of global features while significantly reducing computational overhead.

We have conducted comprehensive evaluations of EMReady2 on three diverse test sets covering different types of maps at 2–10 Å, and compared it with existing methods, including DeepEMhancer, phenix.auto_sharpen, EMReady, and CryoTEN. The evaluation results demonstrate that EMReady2 significantly outperforms existing methods. EMReady2 can enhance the quality of maps for both proteins and nucleic acids. Also, EMReady2 shows effectiveness in enhancing maps across a wide range of resolutions, from 2 to 10 Å. Moreover, EMReady2 can be applied to the maps reconstructed not only by cryo-EM SPA but also by cryo-ET STA. We also demonstrate that EMReady2 improves the map interpretability in terms of *de novo* model building and is robust to map heterogeneity. The runtime analysis confirms its computational efficiency, supporting the processing of larger and more complex maps. It is anticipated that EMReady2 will serve as an enormously valuable tool in the workflow of cryo-EM model building in structure biology.

## 4 Methods

### 4.1 Network architecture

Figure 1a shows the overall architecture of bidirectional Mamba-conv UNet (BiMCUnet). Given an input density map, an initial 3D convolution extracts low-level volumetric features and expands the channel dimension. The encoder contains three downsampling stages, each of which composes of BiMamba-Conv (BiMC) Blocks followed by a convolution that halves the resolution and doubles the channels. At the bottleneck, BiMC Blocks capture contextual information at the coarsest scale. The decoder mirrors the encoder with transposed convolutions and BiMC Blocks, and incorporates skip connections via element-wise addition to preserve spatial detail. Finally, a 3D convolutional tail projects the features to the output domain.

### 4.2 BiMamba block

The BiMC Block is a composite module designed to jointly model local and contextual features in 3D density maps. The input features are split into two branches by a 1*×*1 convolution, which are then processed separately by residual Conv Blocks(RConv) and BiMamba Blocks(BiM). This process can be formulated as follows

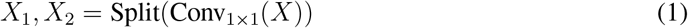

*X*_1_ and *X*_2_ are then fed into a BiM block and a RConv block, respectively, yielding

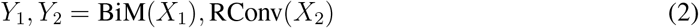

Finally, *Y*_1_ and *Y*_2_ are concatenated as inputs to a 1 *×* 1 convolution, which maintains a residual connection with the input x. The final output of the BiMC block is thus given by

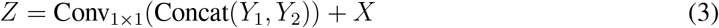

Mamba is based on the structured state space sequence model (S4)^40–43^, which models sequence data as a linear continuous-time dynamical system with paramete **A, B**, and **C**. The continuous system is expressed as follows

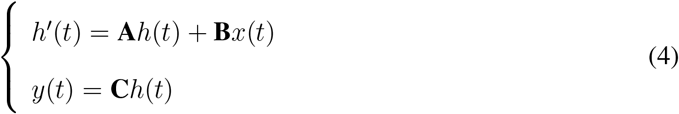

The discrete-time version is obtained by applying a time discretization step **Δ**, converting continuous matrices to discrete counterpart 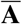 and 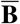 as follows

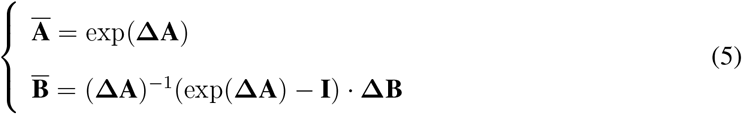

The output is computed via a global convolution kernel 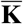 as follows

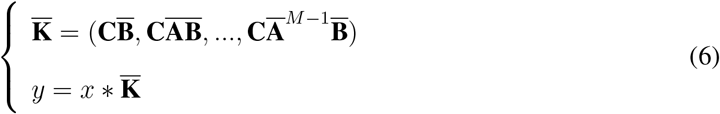

Bidirectional Mamba (BiMamba)^45^ extends the standard Mamba architecture by introducing a dual-branch design, where one branch processes the input sequence in the forward direction, and the other processes the time-reversed sequence in parallel. To integrate information from both directions, the outputs from the two branches are aligned and fused.

Specifically, *y*_*f*_ denote the output from the forward state space model, and *y*_*b*_ denote the output from the backward state space model applied to the reversed input sequence. The backward output is flipped to match the order of the forward output. The two are then combined through element-wise addition, followed by a learned linear projection

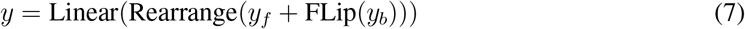

The residual Conv block consists of two consecutive 3D convolutional layers with 3×3×3 kernels, each followed by Filter Response Normalization(FRN). The input feature is added to the output of these layers via a residual connection

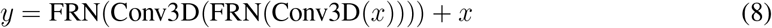

### 4.3 Data collection

In this study, all EM density maps and the corresponding atomic structures in our dataset are downloaded from the EMDB^47^ and the PDB^48^, respectively. To build a consensus and high-quality dataset that covers different types of EM density maps, all maps and PDB structures that meet the following criteria are collected: (i) resolution between 2.0–10.0 Å; (ii) reconstructed by single-particle analysis (SPA) or subtomogram averaging (STA); (iii) a unique correspondence between structure and density map; (iv) without any unknown atom/residue; (v) the total number of *Cα* and *C*4^*′*^ atoms falls within the range of 20–2000; (vi) cross-correlation coefficient calculated by UCSF Chimera^49^ between the structure and the density map is greater than 0.6; (vii) CC_mask calculated by phenix.map_model_cc^50^ between the structure and the density map is greater than 0.5. Further manual inspection is conducted to exclude maps that exhibit obvious inconsistencies with the associated PDB model from our dataset.

To remove the redundancy in our dataset, we use MMseqs2^51^ to cluster the samples in our dataset by using a coverage and a sequence similarity thresholds of 0.8 and 0.3 for proteins and 0.8 and 0.8 for nucleic acids, respectively. The clustering results in a non-redundant set of 549 maps, which are then divided into 348 training samples, 65 validation samples, and 136 testing samples. We further divided the test samples into three distinct sets to address different evaluation needs. The first test set comprises all 118 cryo-EM single-particle analysis (SPA) maps with resolutions ranging from 2.0 Å to 10.0 Å. The second test set includes 18 cryo-electron tomography subtomogram averaging (STA) maps with resolutions of 3.0–10.0 Å. For each map in this set, more than 10% of the residues in the corresponding PDB structure are nucleic acids. The third test set focuses on nucleic acid–containing cryo-EM SPA maps and consists of 18 maps with resolutions between 2.0 Å and 7.0 Å. To evaluate the improvement in map interpretability achieved by EMReady2, we extracted all chains from test-set structures with resolutions better than 5.0 Å, yielding a total of 832 chains, including 753 protein chains and 79 nucleic acid chains.

### 4.4 Generation of local resolution-dependent simulated maps

For each sample in the training set, the corresponding training objective, i.e., simulated map, is generated from its associated PDB structure. Instead of applying a uniform resolution across the entire structure, we convert the PDB structure to the simulated map in a heterogeneity-aware manner, where the local quality is represented by the Q-score^46^. Specifically, for each atom in the PDB structure, the corresponding local resolution 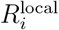 is calculated as follows,

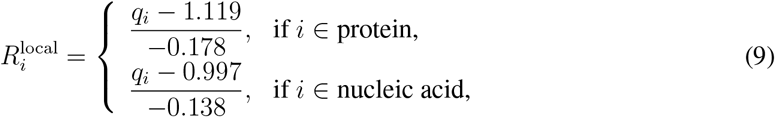

where *q*_*i*_ denotes the Q-score for the *i*-th atom. The final resolution that is used to generate the simulated map from the atom is a combination of global and local resolutions as follows,

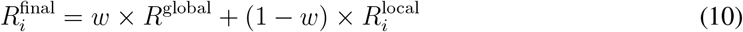

where *w* is set to 0.8 to balance the contributions from global and local resolutions.

Thus, given a PDB structure of *N* atoms, the density value *ρ* on grid point **x** of the simulated map can be calculated as follows,

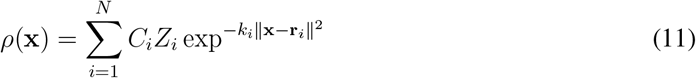

where *Z*_*i*_ and **r**_*i*_ are the atomic number and the position vector of the *i*-th heavy atom (*i* = 1, 2, …, *N*), respectively. *k*_*i*_ and *C*_*i*_ are derived from the final resolution, i.e., 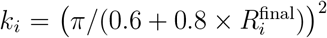 and *C*_*i*_ = (*k*_*i*_*/π*)^1.5^.

### 4.5 Network training

To train the network of EMReady2, we first unify the grid size of all maps in the training set to 1.0 Å through cubic interpolation. Next, the density values of each map are clipped to the range of 0 to the 99.999-percentile of the entire map. Finally, each pair of experimental and simulated maps is segmented into a series of pairs of volume slices using a sliding window of shape 80 *×* 80 *×* 80 and a sliding stride of 48. Non-positive volume slices are excluded in order to ensure effective training.

EMReady2 employs BiMamba-Conv-UNet as its network architecture for post-processing EM density maps. The network consists of three encoder, one bottleneck, and three decoder BiMamba-conv (BiMC) blocks with skip connections. EMReady2 is implemented using PyTorch 2.3.1^52^. Data augmentation is applied during training. Namely, the input pairs of volume slices are augmented by random 90° rotations, followed by randomly cropping a smaller slice of shape 64 *×* 64 *×* 64 from the original slice of shape 80 *×* 80 *×* 80.

The input of our network is density slices of shape 64 *×* 64 *×* 64 with a grid interval of 1.0 Å. The outputs of our network are post-processed density slices of the same size. A combination loss function is adopted by EMReady2 to measure the difference between the output post-processed volume slices and the target simulated slices. This loss function consists of two terms. One is the smooth L1 loss, which is more sensitive to local differences of density values. The other is the structural similarity (SSIM) loss, which reflects the non-local correlation of density slices. Given a post-processed volume slice *Y* ^pred^ and its corresponding simulated volume *Y* ^GT^ of shape *M×M×M* (*M* = 64 in this study), the smooth L1 loss is given as follows,

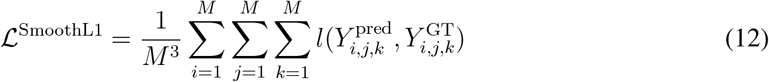

where 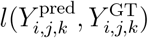 is the smooth L1 distance between the post-processed volume slice and the simulated volume slice at position (*i, j, k*). The smooth L1 distance is defined as follows,

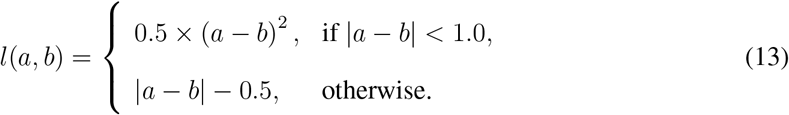

The SSIM Loss measures the contrast and structural similarity between the post-processed and the simulated volume slices, which is given as follows,

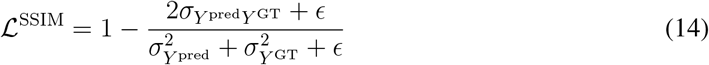

where 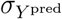 and 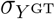 are the standard deviations of the post-processed volume slice and the simulated volume slice, respectively, 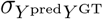 is the covariance between two slices, and *ϵ* = 1*e −* 6 is introduced to prevent division by zero. By combining the smooth L1 loss and the SSIM loss, the final loss function used to train EMReady2 can be expressed as follows,

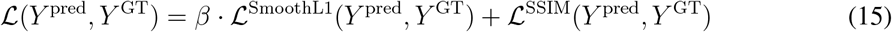

where the weighting factor *β* is empirically set to 100.0 to balance the contributions from local and non-local loss terms.

EMReady2 was trained on four NVIDIA Tesla A100 GPUs with a batch size of 80. The network was optimized using the Adam optimizer with a learning rate of 2 *×* 10^*−*4^, updated via backpropagation. An exponential moving average (EMA) was employed during training to stabilize and enhance model learning. Training was terminated once the loss function had fully converged, typically after approximately 500,000 steps. The choice of hyperparameters for EMReady2 was guided by empirical observation and computational efficiency. Every 400 steps, the model was evaluated on a validation set by computing the mean validation loss, and the model with the lowest validation loss was selected for evaluation.

### 4.6 Evaluations of map quality and interpretability

We conducted comprehensive evaluations of EMReady2 on a total of 136 EM maps. For each density map, we first chunk it into overlapping volume slices of shape 64 *×* 64 *×* 64 with a sliding stride of 24. Subsequently, these overlapping slices are fed into the selected EMReady2 model with the least validation loss. Finally, the resulting slices from the network are re-assembled into the output post-processed map and the overlapping sections are averaged.

Three programs are utilized to measure the quality of maps before and after post-processing. The first is phenix.mtriage^50^, which is used to calculate the FSC-0.5 (the resolution at which the map-model Fourier shell correlation falls to 0.5). The second is the MapQ plugin of UCSF Chimera, which reports Q-score^46^. The Q-score measures the correlation between the density around each atom and a reference Gaussian density function. The main-chain (MC) Q-score refers to the Q-score calculated exclusively for backbone atoms of proteins and nucleic acids. Specifically, we used the *N, Ca, C*, and *O* atoms for proteins, and the *P, O*1*P, O*2*P, O*5^*′*^, *C*5^*′*^, *C*4^*′*^, *C*3^*′*^, and *O*3^*′*^ atoms for nucleic acids. The third tool is phenix.map_model_cc^50^, which reports three different real space correlation coefficients between the map and an ideal map generated from the atomic model: CC_box, CC_mask, CC_peaks. The CC_box is calculated using the map values in the entire map, the CC_mask uses the map values inside a masked region around the atomic model, and the CC_peaks only considers those high-density regions in the map. Among the five metrics, the lower value is the better for FSC-0.5, while the higher value is the better for the others.

In addition to the above map quality metrics, we also assessed the ability of EMReady2 in improving map interpretability. For a fair evaluation, a *de novo* model building tool, phenix.map_to_model^11^ is used to build the atomic models from the map regions segmented within 4.0 Å from each of the protein and nucleic acid chains in our first test set. Residue coverage and sequence recall, calculated by phenix.chain_comparison, are used to assess the accuracy of the built model of protein or nucleic acid relative to the PDB model. Protein residues are represented by *Cα* atoms, while nucleic acid residues are represented by *C*4^*′*^ atoms. Residue coverage is the proportion of residues in the built model that have matched residues (distance within 3.0 Å) in the PDB model. Sequence recall is the fraction of the residues in the built model that have correct residue types compared to those in the PDB structure.

### 4.7 Comparison with related methods

EMReady2 is compared with state-of-the-art methods, including DeepEMhancer^31^, EMReady^33^, phenix.auto_sharpen^27^, and CryoTEN on different test sets. For phenix.auto_sharpen, EMReady and CryoTEN, the default parameters were used. For DeepEMhancer, which provides three different models: ‘tightTarget’, ‘wideTarget’, and ‘highRes’, we reported the evaluation results in a combinatorial manner. Specifically, for cryo-EM SPA maps, we reported the results from the “highRes” model for maps with resolutions higher than 4.0 Å, and the results from the “tightTarget” model for maps with lower resolutions. For cryo-ET STA maps, due to their generally lower quality, we reported the results from the “wideTarget” model. For the evaluation of map interpretability in model building, we reported the results from the best-performing “wideTarget” model.

## Acknowledgments

This work was supported by the National Natural Science Foundation of China (grants No. 32161133002 and 62072199 to S.H., grant No 62276108 to X.W.) and the startup grant of Huazhong University of Science and Technology.

## Author contributions

S.H. conceived the project, S.H., X.W., and J.H. supervised the project. H.C. and Y.Z. implemented the methods and performed the experiments. H.C., T.L., and S.H. analyzed the data. H.C., Y.Z, T.L. and J.C. tested the program. H.C., Y.Z, J.H., and S.H. wrote the manuscript. All authors read and approved the final version of the manuscript.

## Data availability

The raw data of the evaluation results are provided in the Article. All published data sets used in this paper were taken from the EMDB and PDB.

## Code availability

The EMReady2 package is freely available for academic or non-commercial users at https://github.com/huang-laboratory/EMReady2/.

